# Beyond the word and image: III. Neurodynamic properties of the semantic network

**DOI:** 10.1101/767384

**Authors:** Anne-Lise Jouen, Nicolas Cazin, Sullivan Hidot, Carol Madden-Lombardi, Jocelyne Ventre-Dominey, Peter Ford Dominey

## Abstract

Understanding the neural process underlying the comprehension of visual images and sentences remains a major open challenge in cognitive neuroscience. We previously demonstrated with fMRI and DTI that comprehension of visual images and sentences describing human activities recruits a common semantic system. The current research tests the hypothesis that this common semantic system will display similar neural dynamics during processing in these two modalities. To investigate these neural dynamics we recorded EEG from naïve subjects as they saw simple narratives made up of a first visual image depicting a human event, followed by a second that was either a sequentially coherent narrative follow-up, or not, of the first image. In separate blocks of trials the same protocol was presented using sentences. Analysis of the EEG signal revealed common neural dynamics for semantic processing across image and sentence modalities. Late positive ERPs were observed in response to sequential incoherence for sentences and images, consistent with previous studies that examined coherence in these two modalities separately. Analysis of oscillatory power revealed increased gamma-band activity for sequential coherence, again consistent with previous studies showing gamma increases for coherence and matching in sentence and image processing. Multivariate analysis demonstrated that training a classifier on data from one modality (images or sentences) allowed reliable decoding of the sequential coherence of data from trials in the untrained modality, providing further support for a common underlying semantic system for images and sentences. Processing sequential coherence of successive stimuli is associated with neural dynamics that are common to sentence and visual image modalities and that can be decoded across modalities. These results are discussed in the context of EEG signatures of narrative processing and meaning, and more general neural mechanisms for structure processing.

## Introduction

A major function of higher cognitive processing is making sense of the world around us based on accumulated experience that is organized in a narrative structure (Bruner 1990). Binder et al (2009) note that knowledge acquired from experience underlies our ability to understand, and forms the basis of the semantic system. In a meta-study of 120 PET and fMRI studies that access meaning from words, they identified a widely distributed network that suggests that semantic representations tap into sensory, motor, affective and cognitive systems recruited in human experience.

We hypothesized that such a broadly distributed semantic coding is not restricted only to verbal material, but rather that there will be a common network for representing meaning issued from all experience, including verbal and also visual image input. This idea is consistent with early work on language comprehension (Biederman 1981, Friedman 1979, Gernsbacher & Faust 1991, Mandler & Johnson 1976), but the neural underpinnings of such a common network remain to be investigated. The current study represents part three in our series of studies to better understand this network, and to test the hypothesis of a common neural system involved in understanding sentences and images. In our first test of this hypothesis, we determined that a common semantic network would be recruited in the comprehension of visual images, and sentences that depict human events (Jouen et al 2015). fMRI and DTI revealed the spatial organization of this network and aspects of its connectivity. Interestingly, the common network was quite similar to that identified by Binder et al (2009), including major activation in the angular gyrus and temporo-parietal cortex. This was inspired by the groundbreaking work of Vandenberghe et al (1996) in understanding the common semantic system, and extended their approach from simple images and single words, to rich images and full sentences.

In our second investigation of this hypothesis we performed detailed mapping of the white matter pathways and functional connectivity linking the cortical nodes of this distributed network, with major hubs in the anterior temporal cortex and the temporo-parietal cortex (Jouen et al 2018). The dense connectivity of these areas highlights their roles in integrative processes. Such densely connected temporo-parietal regions could serve as anchor points for the convergence of multimodal cortical representations during experience. During comprehension, activation of such convergence zones could then allow for divergent reactivation of multimodal areas in reconstructing meaning (Lallee & Dominey 2013, Meyer & Damasio 2009). While fMRI and DTI thus provide a view of the spatially distributed network organization of the semantic system and its connectivity that will be engaged in making meaning from narrative, a different approach is required for characterization of the temporal dynamics of this integrative processing. Given its temporal precision, electroencephalography (EEG) is a more suitable tool for investigating the temporal unfolding of processes in narrative integration. We thus set out to compare the temporal dynamics of the EEG signal in visual image and sentence processing using short (two-element) narratives that allowed us to manipulate the sequential coherence of the successive stimuli within the narratives. The goal is to determine if there are EEG responses that are common to image and sentence processing, which would provide further evidence for common underlying neural mechanisms.

Written and visual narratives have been investigated separately using EEG in protocols where different dimensions of narrative coherence have been manipulated. Late ERP positivities with some variability in their localization and timing are frequently observed in these coherence manipulations. This can be seen during violation of the expectation or goal in visual narrative (Cohn et al 2014, Sitnikova et al 2008), and violation of semantic expectations built up earlier in a text in verbal narrative (Bornkessel-Schlesewsky & Schlesewsky 2008, Brouwer et al 2012, Paczynski & Kuperberg 2012, Xiang & Kuperberg 2015). While there are a variety of experimental manipulations that can modulate late positivities, there is converging evidence that these responses can be associated with aspects of semantic incoherence (Brouwer et al 2012). These results from narrative meaning processing using verbal and visual image stimuli indicate that encountering unfulfilled expectations or predictions can invoke on-line processes that are often revealed by late ERP positivities.

In addition to these late positivities, other ERP responses have been associated with different aspects of semantic processing, including earlier positivities in the 300s time-frame (e.g. Polich 2007), and discourse-related negativities in the 400ms time frame (Hagoort & van Berkum 2007). The timing and distribution of these effects can be modulated by different dimensions of the stimuli. For our purposes, what is important is the identification of ERP responses that can reliably reveal (or not) common processes for sentence and image processing.

Such aspects of cognitive processing can also be characterized in terms of oscillatory power in neuronal activity. EEG gamma-band activity has been associated with semantic processing in language (Hagoort et al 2004, Hald et al 2006) and in multisensory semantic matching (Schneider et al 2008). The gamma-band activity appears to increase in situations of semantic congruence. Thus, gamma band responses were observed to increase in responses to semantically correct vs. violation sentences (Hald et al 2006). Likewise, gamma-band responses increased in response to semantically matching vs mismatching stimuli (Schneider et al 2008) in a multisensory semantic priming task using visual and auditory stimuli. We can thus predict that gamma-band responses would be increased in conditions of increased coherence between successive stimuli both for sentences and images.

While the demonstration of related ERP and time-frequency effects for image and sentence processing suggest a common underlying mechanism, demonstration of cross-modal decoding (i.e. training a decoder on image data and decoding sentence data, and vice-versa) would provide more convincing evidence that the neural dynamics in the two cases share a common component. As stated by Shinkareva et al (2011) successful discrimination in one stimulus format based on neural activation patterns elicited by a different stimulus format implies consistent triggering of semantic representations by these different stimulus formats. Thus the neural dynamics represented in the decoder represent high level properties, and not just perceptual properties associated with picture or sentence stimulus formats (Shinkareva et al 2011). Motivated by this approach we thus consider such a cross-modal decoding in the current research, based on the EEG signal.

The EEG signal is sensitive to diverse cognitive functions and has been used to decode cognitive states. For example, attentional focus in an auditory “cocktail party” context can be decoded from single trial EEG (O’sullivan et al 2014), and EEG has been used to discriminate which one of seven different musical fragments a subject is listening to (Schaefer et al 2011). This type of decoding should be suitable to test our hypothesis of the existence of common neurophysiological processes for the semantic processing of visual and written narratives. If there are common neural processes involved in the understanding of sentences and images, they should display a common spatial and temporal distribution of comprehension-related activity.

We focus here on the detection of a form of narrative coherence in the sentence and image domains. We adapt the protocol from Jouen et al (2015) to the EEG domain, retaining the overall structure of the protocol that involved the presentation of natural visual images depicting human activity, and in separate blocks of trials, short whole sentences describing the same type of human activity. We introduce an aspect of temporal succession corresponding to simple narratives. In both the sentence and the image domains, two successive stimuli are presented, separated by a variable delay. The second stimulus can be sequentially coherent or incoherent in a narrative context with respect to the first. Brain responses to these stimuli are analysed to determine if there is evidence for a common processing of sequential coherence in sentence and image modalities, in the form of common EEG responses to incoherence across these modalities.

## Methods

### Participants

Eighteen healthy right-handed volunteers participated in the experiment (10 females, 8 males, native French speakers, without prior neurological history, 25.5 ± 4.1 years). The study was conducted in accordance with the Declaration of Helsinki, and all participants were advised of the physical details of the experiment, and gave their informed written consent to participate in the experiment.

**Figure 1.**
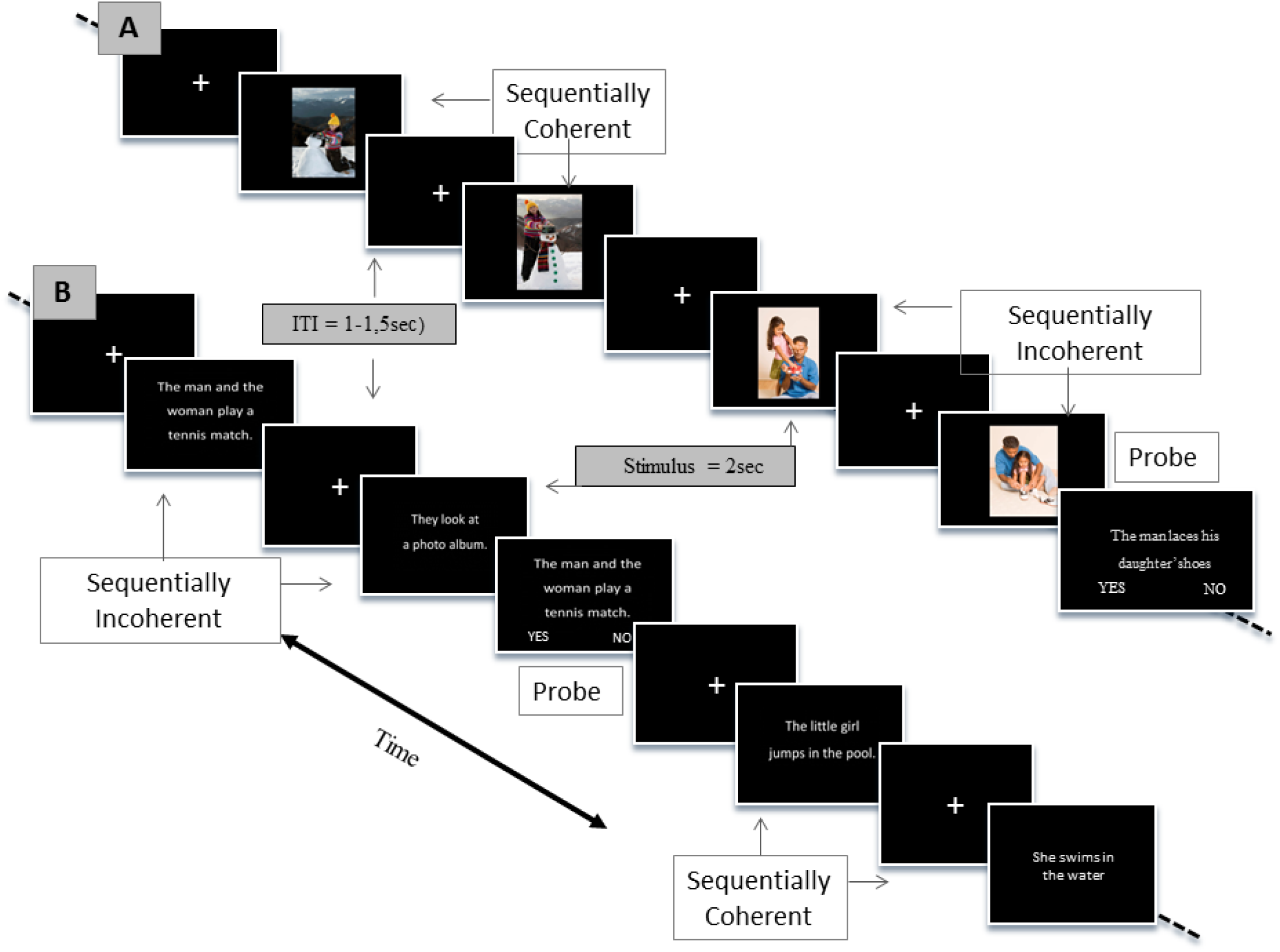
Temporal unfolding of the experimental protocol for the image (A) and sentence (B) conditions.

### Stimuli

The temporal unfolding of the paradigm is presented in Figure 1. For the image conditions one-hundred and twenty (120) image pairs were selected from the Getty photo database (http://www.gettyimages.fr/). A set of sixty pairs of images made up sequentially coherent narratives, and a second set of sixty pairs of images made up sequentially incoherent narratives. When a narrative was sequentially coherent, the images depicted the same people performing a logical sequence of activities (e.g. entering a bakery; buying bread). Sequentially incoherent pairs depicted the same people performing two unrelated activities. Likewise, for the sentence conditions, sixty pairs of sentences were created that made up sequentially coherent narratives, and sixty pairs for incoherent narratives. In the same way that images can implicate the same people in different situations over the presentation of two successive stimuli, the sentences describing people performing an action can follow-up (or not) in the second sentence. To reproduce a similar effect as with the images, that is, that certain information does not need to be re-analysed (e.g. the identity of the people, their different roles in the relations that link them) we chose to use complete nominal specification for stimulus 1 (e.g. “The man and the woman looked at the movie announcements”) and replace the agents with pronouns to refer to the same protagonists in stimulus 2 (e.g. “<u>They</u> bought tickets at the counter.”). This is consistent with pragmatic discourse rules and avoids the repeated name penalty (Gordon et al 1993). Example sentences and images are illustrated in Figure 1. The images were controlled and counterbalanced for the number of people illustrated (1 or 2). Additional sample sentences are:

Sequentially Coherent:

S1- The man opens the driver side door. S2 - He starts the car.
Sequentially Incoherent:

S1 - The woman lounges by the pool. S2 - She irons the laundry.

### Experimental Paradigm

Subjects were seated in front of a visual display. Visual stimuli (sentences or images) were presented at the center of the screen, subtending a visual angle of approximately 5°. Subjects saw a visual image (or read a sentence) depicting a human event, and after a pause saw a second image (or sentence) that was either sequentially coherent, or not, with the first image (or sentence). Stimulus one and two were always in the same modality (sentence or visual image).

A trial started with a fixation point for a variable delay of 0.4-1 sec., then the first stimulus was presented for 2 sec and after a delay of 1-1.5 seconds the second stimulus was presented for 2 sec followed either by a fixation point or a question for 1-1.5 sec. Two thirds of the trials were followed by probe questions in order to maintain vigilance. Subjects responded to the question by yes or no by pressing with their right hand respectively a right and left key on a button pad. The probe question was always related to the stimuli of the previous trial to avoid any memory confounds. Trials were blocked by modality (image or sentence), two blocks per modality, for a total of 4 blocks. Each block had 30 coherent, and 30 incoherent trials.

### EEG acquisition and preprocessing

We acquired continuous neural activity with 64 channel EEG (Biosemi, ActiveTwo, version 5.36) sampled at 2Khz while subjects performed the task. EEG data was processed using EEGLAB. Preprocessing was performed with the FASTER plugin for EEGLAB (Nolan, Whelan et al. 2010), with bandpass filter from 1 Hz to 95 Hz, notch filter (50 Hz), artifact rejection and epoching from −0.5 to 3 s relative to the stimulus onset. All electrodes were referenced with respect to the mastoids. Artifacts from eye movements, blinks and temporal muscle activity were identified and removed using a second-order blind identification (SOBI) algorithm implemented in the EEGLAB toolbox interface (Lio and Boulinguez 2013).

### EEG Analysis

The EEG signal was analysed in terms of task related event-related potentials (ERPs), time frequency analysis, and cross-modal (sentence vs. image) decoding approaches.

a. ERP analysis: Average waveforms were computed across all trials per condition, using a baseline subtraction with the interval [-200 0 ms]. Repeated measures ANOVAs and Wilcoxon non-parametric tests were performed using MATLAB, on the factors: Modality (Image, Sentence), Sequential Coherence (Sequential, Non-Sequential) and Order (First, Second stimulus) for different time windows. Holm’s method was used to correct for multiple comparisons (Holm 1979). In order to be more complete, the analysis was performed for multiple extended time windows including 250-350ms, 500-625ms, 750-1000ms, 1600-1800ms windows. These windows were chosen as a function of classic stimulus driven responses and the morphology of the ERP response in our recording. The 250-350ms response covers the classic stimulus driven P2 and context-processing P3 (Polich 2007). The 500-625 overlaps with a late negativity identified for semantic and mental imagery processing (West & Holcomb 2000). The 750-1000ms overlaps with late positivity effects observed in sentence and discourse processing e.g. (Brouwer et al 2012, Kuperberg 2007). The 1600-1800ms corresponds a previously little explored period where we observe task related effects.
b. Near Gamma Band (NGB 35-45 Hz) Time-frequency analysis: Time-frequency (TF) event-related spectral perturbation (ERSP) representations were obtained by computing the mean squared norm of the convolution of complex Morlet wavelets with the EEG data, across the trials per condition. We used wavelets with a 3-cycle width and frequencies ranging from 1 to 80 Hz, at 1 Hz intervals. We performed a single-trial baseline correction, dividing the ERSP power at each time point by the average spectral power in the pre-stimulus baseline period at the same frequency before averaging and taking a log-transformed measure of the percentage of ERSP as described in Grandchamp and Delorme (2011). Repeated measures ANOVAs and post-hoc comparisons with Greenhouse-Geisser correction for sphericity were performed using MATLAB, on the factors: Modality (Image, Sentence), Sequential Coherence (Sequential, Non-Sequential) and Order (First, Second stimulus) on the ERSP in the near-gamma band (35-45 Hz) for sliding time windows of 200 ms. Holm’s method was used to correct for multiple comparisons (Holm 1979).
c. Cross-modal decoding: The objective of this analysis is to determine if multivariate discrimination of neural activity patterns during sequential coherence processing in one stimulus format can be used to perform the same discrimination in the other stimulus format. This would argue that the neural activity captured by the decoder reflects semantic property representation, and not just perceptual features associated with the stimulus formats (Shinkareva et al 2011).

To achieve this, principal component analysis (PCA) was performed, using MATLAB, on the signals from the 64 electrodes over all subjects and all conditions of sequential coherence (Sequential, Non-Sequential), modalities (Image, Sentence) and stimulus order (first and second) confounded, starting from −0.5 to 3 s relative to the stimulus onset. Data from sequential vs non-sequential, image vs. sentence, and first vs. second stimulus trials was then separated and analyzed using the resulting principal components, to determine if the principal components were sensitive to these task factors, particularly the sequential coherence.

After establishing the relation between task variables, including sequential coherence, and the principal components, we then performed a cross-modal decoding. That is, we performed PCA on data from image processing and determined whether the resulting principal components could be used to discriminate sequential coherence in the sentence data (and vice versa).

**Figure 2.**
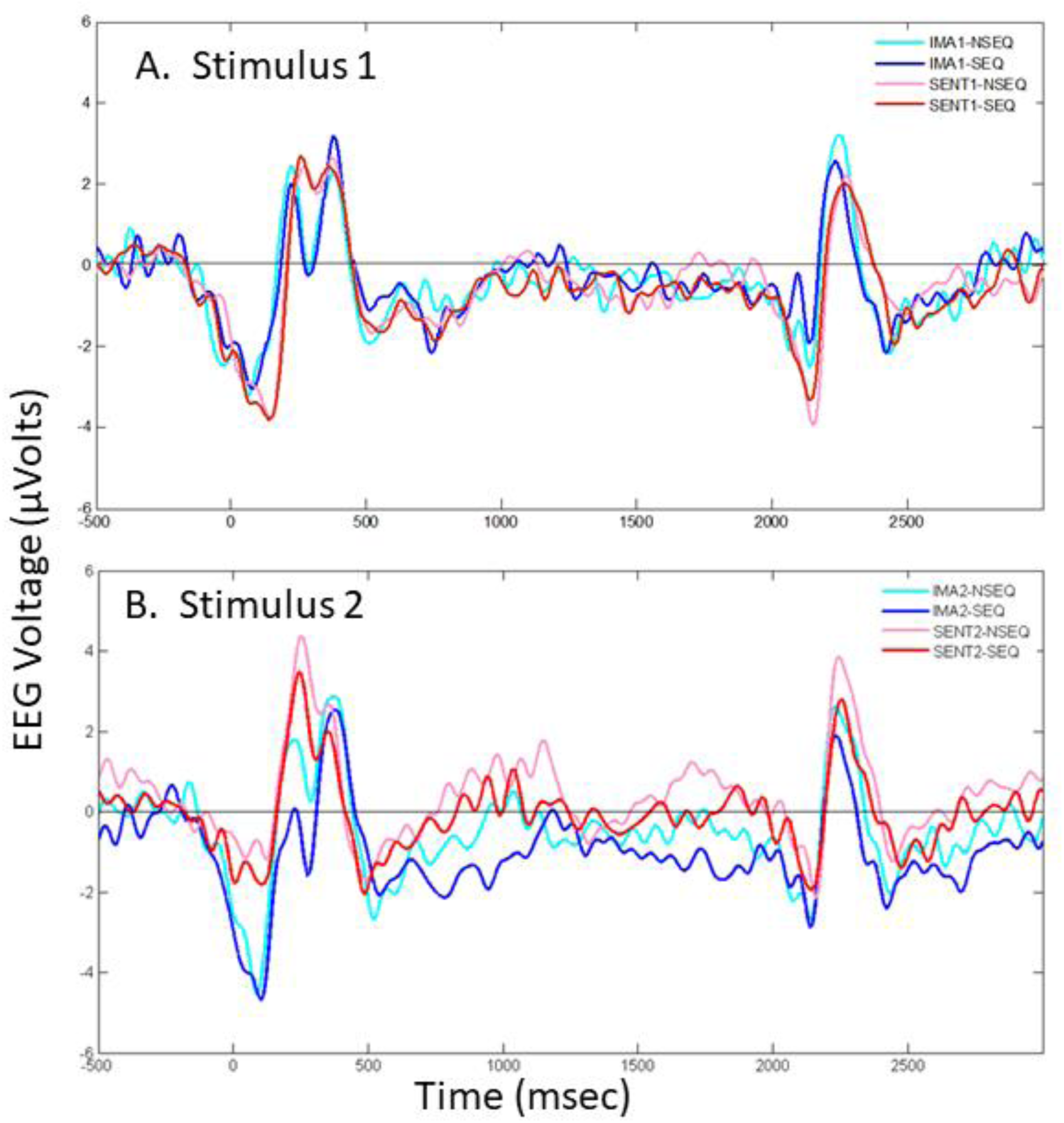
Comparison of ERP responses to first and second stimuli. The large deflections seen at the start and end of the analysis epoch reflect visual onset and offset responses, respectively. A. Stimulus1 – small effects of modality are seen, but no effects of sequential coherence are visible (as expected). B. Stimulus 2 - Sequential coherence effects become visible. IMA1 – image 1; SENT1 – sentence 1; (same for 2); NSEQ – non-sequential; SEQ – sequential.

## Results

### EEG Analysis

Figure 2 illustrates an example of the effects Order (First, Second stimulus) on ERP responses to Modality (Image, Sentence) and Sequential Coherence (Sequential, Non-Sequential) in a representative frontal central electrode FC1. Note that during extended periods during the stimulus presentation (e.g. 750-1000 ms) the modality and sequential coherence effects are absent for stimulus 1, and then become pronounced for stimulus 2. This is supported by the ANOVA. During the 750-1000 ms window there are significant effects for Order [F(1,17) = 6.0, p < 0.05], Modality [F(1,17) = 24.4, p < 0.001], and Sequential Coherence [F(1,17) = 10.8, p < 0.01]. Interestingly there is an Order * Modality interaction [F(1,17) = 11.83, p < 0.01], and an Order * Sequential Coherence interaction [F(1,17) = 16.3, p < 0.001]. Overall, all 64/64 electrodes demonstrated a significant main effect for Modality, 55/64 for Stimulus Order, and 31 for Sequential Coherence. The Modality * Order interaction was observed in 45/64, and the Sequential Coherence * Order interaction in 24/64 electrodes. This indicates rich effects that are highly influenced by the factors manipulated in the task, and that the effects of coherence emerge, as expected only with the second stimulus. We now provide a more specific analysis of these effects. ERP results are presented for four time periods 250-350ms, 500-625ms, 750-1000ms, and a later positivity at 1600-1800ms.

#### Specific Task-Related Effects

Recall that the coherence of a given pair of stimuli can only be discerned by the subject after the presentation of stimulus 2, thus there should be no coherence distinction with respect to the first stimulus, since it must be compared to the second. In order to test for this effect, for each time period we first performed an ANOVA and identified electrodes that displayed a Sequential Coherence * Order interaction. This interaction could reflect a significant Coherence effect only for the second stimulus. For electrodes with a significant interaction we then used non-parametric Wilcoxon signed rank tests to identify those that had a significant Sequential Coherence effect for Images and Sentences for the Second stimulus. Finally we tested whether there was a significant Sequential Coherence effect for Images and Sentences for the First stimulus, which should be absent. Indeed, in none of these cases was this effect found for the First stimulus.

**Figure 3.**
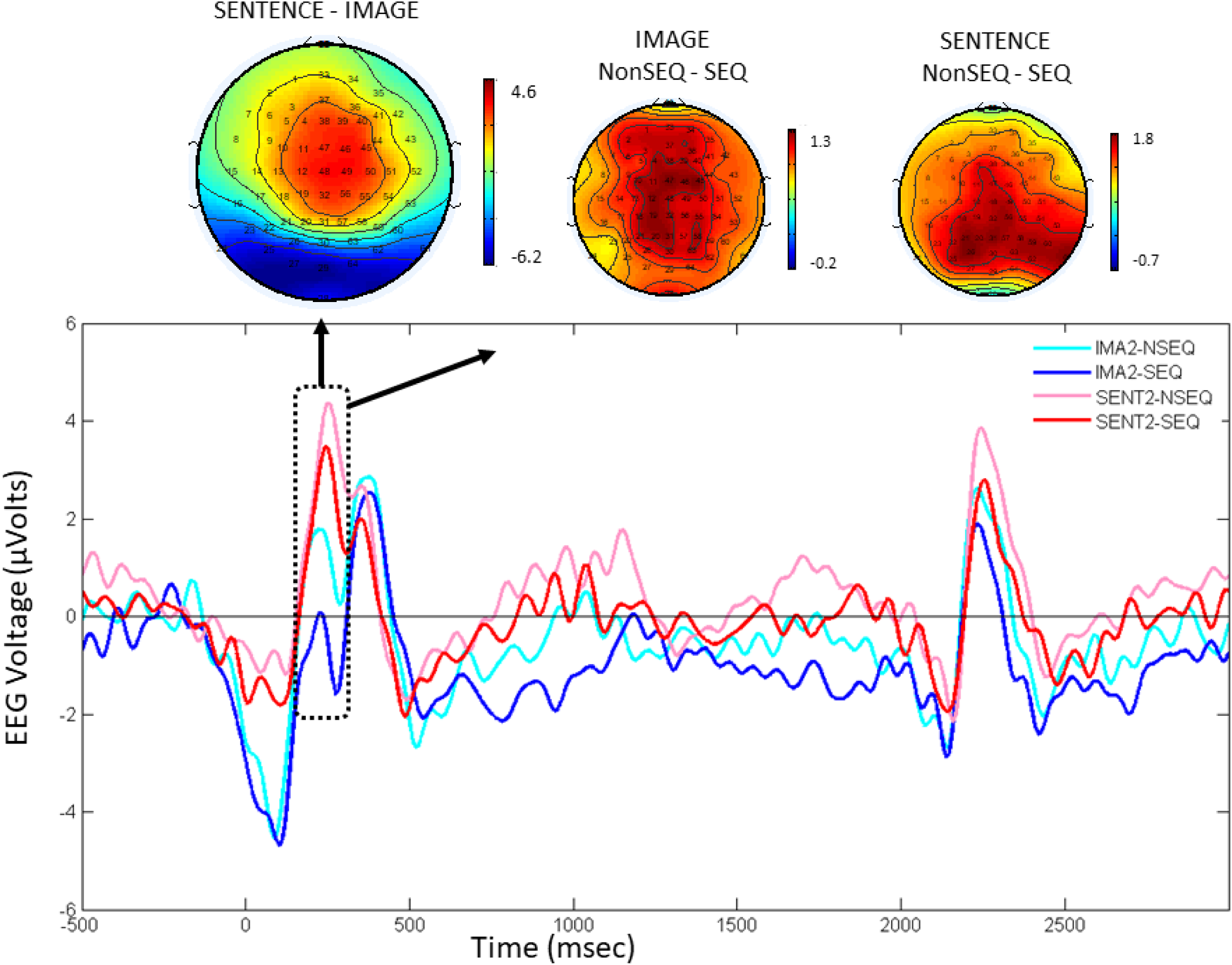
Early Positvity 250-350ms effect. This positivity makes a modality distinction, with a greater positive effect for sentences. It also makes the NonSEQ (sequentially incoherent) vs. SEQ (sequentially coherent) distinction, with an increased positivity for the sequentially incoherent stimuli. Illustrated with Electrode FC1 (11). All values in mico-volts.

**Table 1.** Discrimination of modality and sequential coherence by an Early ERP positivity. Statistical p-values for Wilcoxon signed rank comparisons. Right two columns (Coherence Discrimination) illustrate electrodes with significant coherence differences for sentences and images in the Early Positivity (250 – 350 ms) window. Left two columns (Modality Discrimination) illustrate how in the same window several of these electrodes distinguish sentence and image. All of these electrodes display a significant sequential coherence x order interaction in the ANOVA, reflecting the coherence effect being present only for the stimuli. This demonstrates a form of mixed selectivity (Rigotti et al 2013). second

| Electrode | Modality Discrimination |  | Coherence Discrimination |  |
| --- | --- | --- | --- | --- |
|  | Coherent | Incoherent | Image | Sentence |
| **Central** (C1-12) | 0,005 | 0,025 | 0,0084 | 0,0074 |
| **Centro-Frontal** (FCz-47) | 0,006 | 0,008 | 0.0074 | 0.0096 |
| **Centro-Parietal**<br>(CPz-32) | - | - | 0.0084 | 0.0057 |
| (CP1-19) | - | - | 0.0096 | 0.0043 |
| (CP2-56) | 0,01 | 0,043 | 0.005 | 0.0038 |
| **Parietal** (Pz-31) | - | - | 0.0074 | 0.0043 |
| **Temporal** (T8-52) | - | - | 0.0074 | 0.0096 |

#### 250-350ms Effects

Surface maps and ERP plots in Figure 3 display the modality contrast (sentences vs. images), and the sequential coherence contrasts (NonSEQ – SEQ) for both images and sentences, in the 250-350 timeframe. Interestingly, there is a clear modality effect (discriminating sentences vs images) as well as a sequential coherence effect for sentences and images, as an extended group of frontal central electrodes dispaly both coherence and modality discriminations in this timeframe. These observations are confirmed by ANOVA and Wilcoxon tests. As detailed in Table 1, seven central electrodes displayed the Sequential Coherence * Order interaction, and further displayed a reliable discrimination of NonSEQ – SEQ at p < 0.01 for Stim 2 for Sentences and Images. None of these electrodes had a reliable discrimination of NonSEQ – SEQ for Stim 1. Three of these electrodes also displayed a main effect for modality (see Table 1). Illustrative frontal central electrode C1-12 displays significant effects for Modality [F(1,17) = 9.4, p < 0.01], and Sequential Coherence [F(1,17) = 16.6, p < 0.001], and a significant Sequential Coherence * Order interaction [F(1,17) = 12.5, p < 0.01]. Thus, an early positivity displays a clear coherence effect dependent on whether the subject is exposed to the first or second stimulus, i.e. only for the critical second stimulus, both for sentence and images. This supports the hypothesis of common neurodynamics underlying semantic processing of sentences and images.

**Figure 4.**
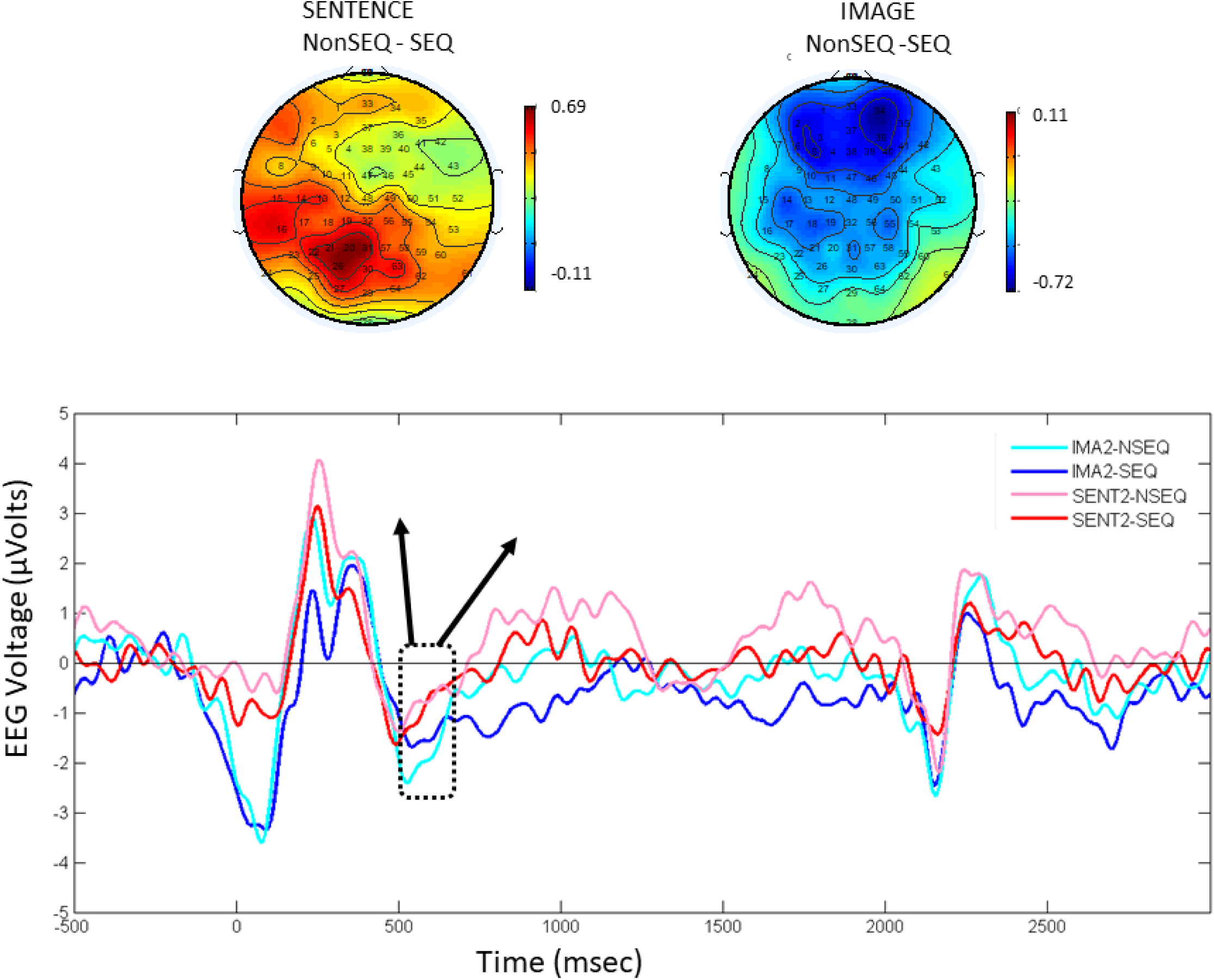
Late Negativity timeframe (525-600ms) there is a significant negativity for NonSEQ (sequentially incoherent) vs. SEQ (sequentially coherent) for images but not sentences. Electrodes 2[AF7], 3[AF3], 6[F5], 36[AF4], 41[F6] (average of these four electrodes illustrated in this figure) display a significant effect, only for the second stimulus.

#### 500-625ms Effects

Surface maps and ERP plots presented in Figure 4 display the sequential coherence contrast for sentences and images in the 500-625ms time frame. We can observe that a sequential coherence discrimination with a frontal negativity is made for the second stimulus, in a number of centrally located electrodes, but only for the Image and not the Sentence modality. Six electrodes displayed the Sequential Coherence * Order interaction, and further displayed a reliable discrimination of NonSEQ – SEQ at p < 0.01 for Stim 2, but only for images. None had an effect for Stim 1. (AF7-2, AF3-3, F3-5, F5-6, AF4-36, F6-41) Illustrative frontal electrode AF7-2 displays a significant effect for Modality [F(1,17) = 7.0, p < 0.01], and a significant Sequential Coherence * Order interaction [F(1,17) = 5.25, p < 0.01], only for images. Thus, during this period a sensitivity to the coherence effect is observed, but only for images.

**Figure 5.**
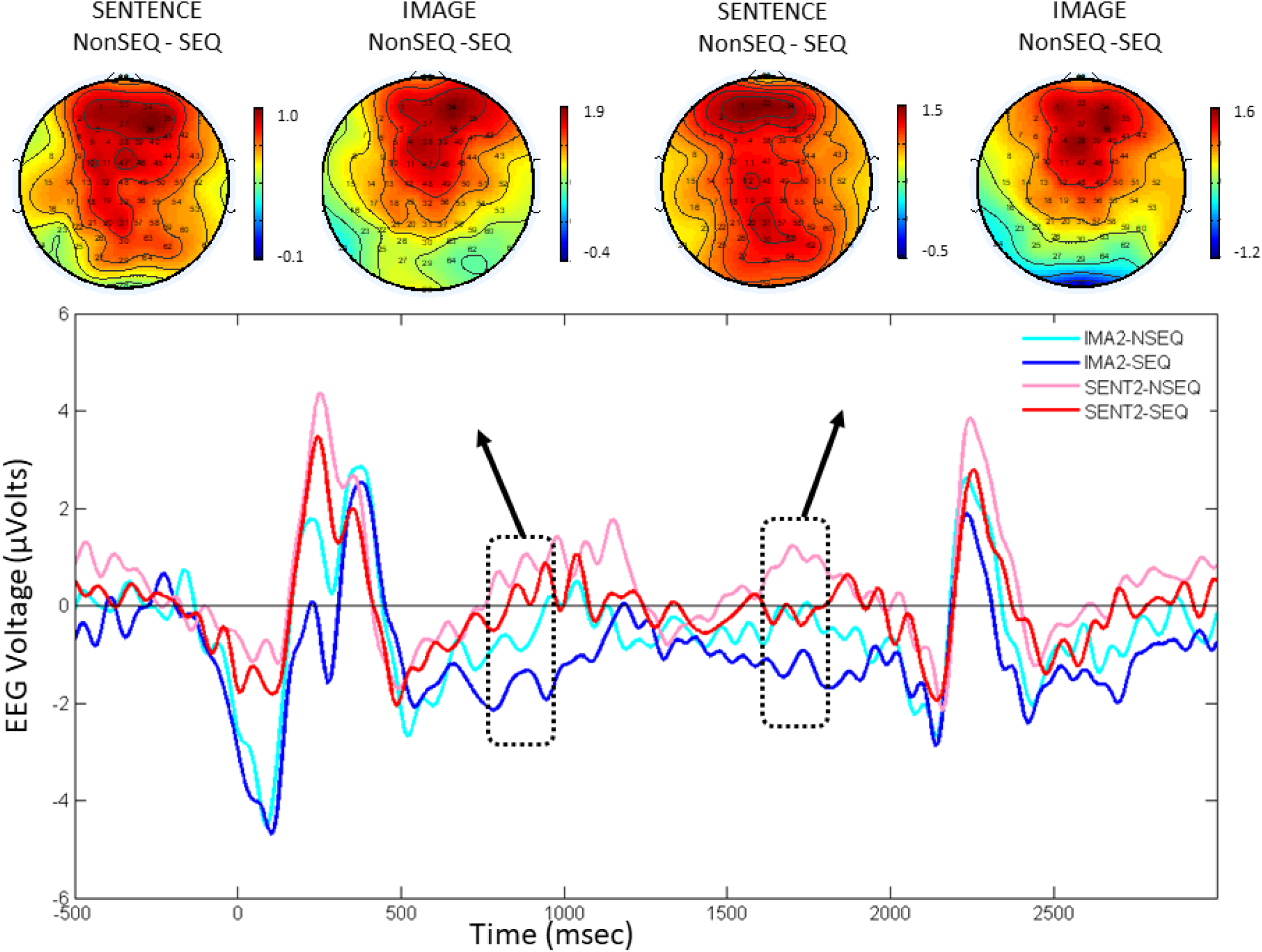
Late positivities (750-1000, and 1600-1800 ms) are sensitive to the NonSEQ (sequentially incoherent) vs. SEQ (sequentially coherent) distinction for sentences and images. Scalp maps for sentences and images NonSEQ-SEQ contrast for 750-1000ms displayed on left, and for 1600-1800 displayed on right. Illustrated with Electrode FC1 (11).

**Table 2.** Discrimination of sequential coherence by Late ERP Positivity. Right two columns (Coherence Discrimination) illustrate electrodes displaying significant sequential coherence differences for sentences and images in the 750-1000 ms window, and the corresponding statistical p-values for Wilcoxon signed rank comparisons. All electrodes except FT8 displayed a significant sequential coherence interaction. A significant effect in the late positivity 1600-1800ms time window was observed in all of these electrodes with the exception of FT8-43, and CP2-56. We also illustrate (Modality Discrimination) how in the same window these electrodes distinguish mixed selectivity. Sentence and image. This demonstrates a form of

| Electrode | Modality Discrimination |  | Coherence Discrimination |  |
| --- | --- | --- | --- | --- |
|  | Coherent | Incoherent | Image | Sentence |
| **Central** (C4-50) | 0.0018 | 0.0012 | 0.0084 | 0.0475 |
| **Fronto-Central**<br>(FC1-11) | 0.0018 | 0.0065 | 0.0014 | 0.0475 |
| (FC2-46) | 0.0018 | 0.0025 | 0.0009 | 0.0386 |
| (FC3-10) | 0.0033 | 0.0084 | 0.0043 | 0.0311 |
| (FC4-45) | 0.0029 | 0.0021 | 0.0016 | 0.0475 |
| (FC6-44) | 0.0043 | 0.0007 | 0.0025 | 0.0429 |
| **Centro-Parietal**<br>(CP2-56) | 0.0002 | 0.0009 | 0.0156 | 0.0386 |
| **Fronto-Temporal**<br>(FT8-43) | 0.0279 | 0.005 | 0.0347 | 0.0429 |

#### 750–1000 ms and 1600–1800 ms Effects

Figure 5 illustrates ERP responses and surface maps for the sequential coherence contrast for sentences and images in the 750–1000 and 1600–1800 ms time frames. We can see similar frontal positivities for the Image and Sentence modalities for both of the late time windows. These observations are confirmed by ANOVA and Wilcoxon tests. Multiple spatially contiguous electrodes displayed the Sequential Coherence * Order interaction, and further displayed a reliable discrimination of NonSEQ – SEQ at p < 0.01 for Stim 2 for Sentences and Images. None of these electrodes had a reliable discrimination of NonSEQ – SEQ effect for Stim 1 (see Table 2). Illustrative frontal central electrode FC1 displays significant effects for Modality [F(1,17) = 24.4, p < 0.001], Sequential Coherence [F(1,17) = 10.8, p < 0.01], and Order [F(1,17) = 6.0, p < 0.05], and a significant Sequential Coherence * Order interaction [F(1,17) = 12.5, p < 0.01]. This interaction reflects the observation that the response to sequential incoherence is only observed when the subject is exposed to the second stimulus. Wilcoxon tests confirmed that only for stimulus 2 there was a significant sequential coherence contrast for both modalities (see Table 2). These two late positivities thus reliably distinguish the sequential vs non sequential coherence contrast for the second stimuli in both sentence and image modalities. Interestingly, as indicated in Table 2, they also make the modality distinction. These late positive effects contribute to the hypothesis that there exist common neurodynamics underlying the processing of semantic coherence in sentences and images.

**Figure 6.**
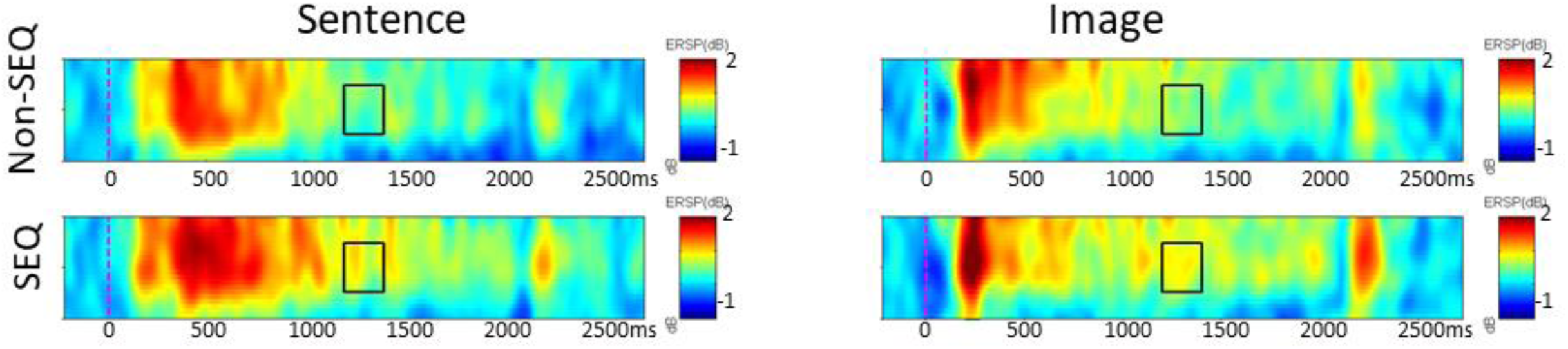
Near Gamma (Hz) oscillations are increased for Stim 2 in the sequentially coherent condition. Panels A and B illustrate TF data for electrode FC5-9 in response to Stim 2 for Sentences in the Non-Sequential and Sequential conditions in the 30-50 Hz range. Marked region in the 1200-1400ms 35-45Hz time-frequency window reveals increased event-related spectral perturbation (ERSP) for the Sequential vs. Non-Sequential conditions. Panels C and D illustrate TF data in response to Stim 2 for Images in the Non-Sequential and Sequential conditions in the 30-50 Hz range. Marked region in the 1200-1400ms 35-45Hz time-frequency window reveals increase ERSP for the Sequential vs. Non-Sequential conditions. For sentences and images there is increased near gamma activity in the 1200-1400ms time-frequency window.

**Table 3.**
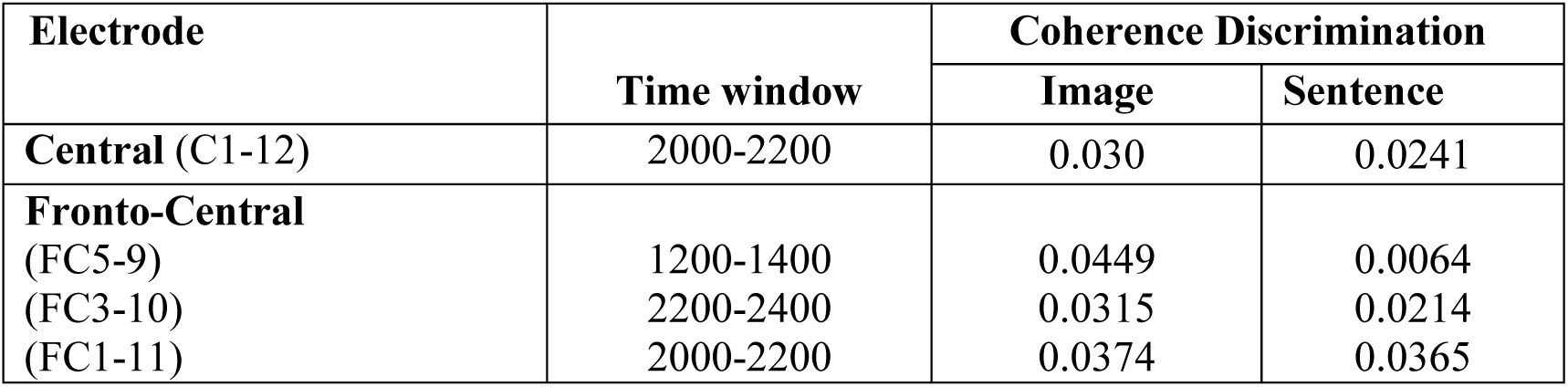
Discrimination of sequential coherence by Near-Gamma oscillatory power. Right two columns (Coherence Discrimination) illustrate electrodes displaying significant sequential coherence differences for sentences and images in the near-gamma (34-45 Hz) range, and the corresponding statistical p-values for Greenhouse-Geisser corrected post-hoc comparisons.

### Near-Gamma Time-Frequency Analysis

Figure 6 illustrates ERSP responses for sentences and images in the sequentially coherent and non-coherent conditions. In the period 1200-1400ms, near-gamma activity (35-45 Hz) appears to be greater for sequential vs non-sequential stimuli, only for the second stimulus. These observations are confirmed by ANOVA and Wilcoxon tests. Multiple left frontal electrodes displayed the Sequential Coherence * Stimulus order interaction, with a significant effect both for images and sentences for the second stimulus and no sequential coherence effect for the first stimulus (see Table 3). In the period 1200-1400ms illustrative frontal electrode FC5-9 displays significant effects for Modality [F(1,17) = 5.3, p < 0.05], and Sequential Coherence [F(1,17) = 4.6, p < 0.05], a significant Modality * Stimulus Order interaction [F(1,17) = 13.0, p < 0.01], and a significant Sequential Coherence * Stimulus Order interaction [F(1,17) = 9.1, p < 0.01]. This interaction means that the sequential coherence response depends on the stimulus order. Wilcoxon tests reveal that in all cases, near-gamma power increases in the sequentially coherent condition for stimulus 2. Summary statistics for all significant effects are provided in Table 3. This effect for sequential coherence for sentences and images contributes to the argument for shared neurodymamics underlying the processing of sentences and images.

**Figure 7.**
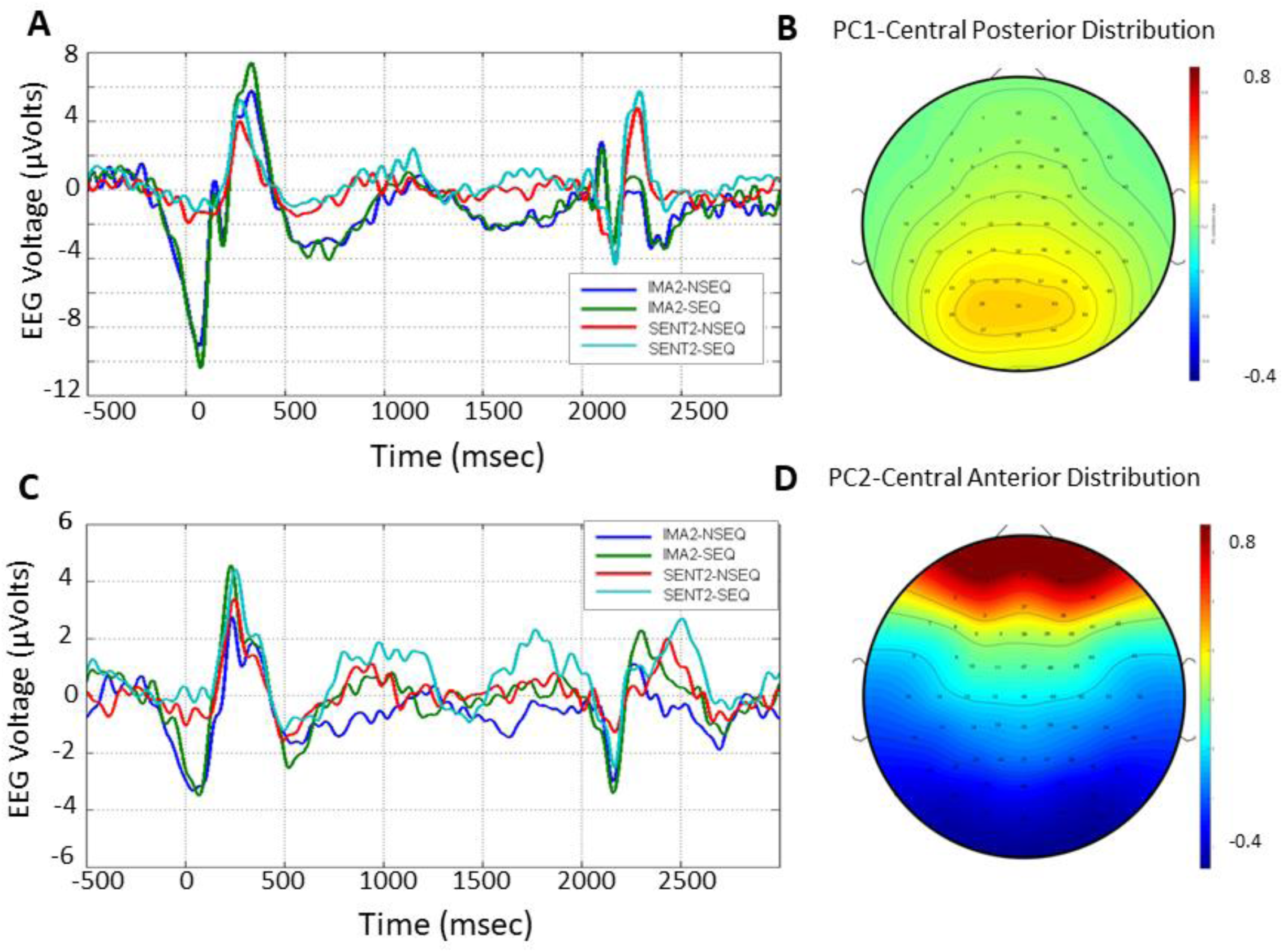
A. Responses of electrodes contributing to Principal Component (PC) 1 to stimulus 2 in all four conditions (IMA,SENT x SEQ,Non-SEQ). Clear separation of IMA vs SENT trials in multiple time windows. B. Spatial distributions of contributions of electrodes to PC1. C. Responses of electrodes contributing to PC2 to stimulus 2 in all four conditions (IMA,SENT × SEQ,Non-SEQ). Clear separation of SEQ and Non-SEQ trials in the 750-1000 and 1600-1800 timeframes. D. Spatial distributions of contributions of electrodes to PC2.

**Table 4.** Sequential coherence discrimination for Stimulus 2. Wilcoxon p values for NonSEQ – SEQ differences for the second stimulus, based on projection of SENT and IMA data for SEQ and NonSEQ trials onto the first two PCs generated by applying PCA to the entire data set. We can observe that between PC1 and PC2 the NonSEQ – SEQ distinction reliably be made.

|  | **Late 1 (750-1000)** |  | **Late 2 (1600 – 1800)** |  |
| --- | --- | --- | --- | --- |
|  | **SENT** | **IMA** | **SENT** | **IMA** |
| **PC1** | 0.159 | 0.200 | 0.005* | 0.159 |
| **PC2** | 0.132 | 0.015* | 0.141 | 0.017* |

### Multivariate Cross-modal decoding

In order to determine the principal components accounting for variability in the EEG signal, we first performed PCA on data from sentences and images, across all subjects and trials. As illustrated in Figure 7, the first two components of the PCA appear to represent the two predominant dimensions in the data which are, respectively, the sentences vs. pictures Modality of the stimuli (Fig. 7A-B), and the Sequential Coherence of stimulus 2 (Fig 7C-D). Wilcoxon tests on the sequential coherence differences reveal that PC1 reliably makes the coherence distinction for Sentence 2 in the second late period (1600-1800ms), and PC2 reliably makes the coherence distinction for Image 2 in the first (750-1000ms) and second late periods (see Table 4). In all comparisons the coherence distinction is never significant for the first stimuli. This indicates that the PCA is sensitive to the distinction between coherent and incoherent stimuli for the key second stimulus.

#### Cross-Modal Validation

A new PCA was performed only on data from image trials, and then the resulting principal components were used to decode the sequential coherence using data from sentence trials. The symmetrical decoding operation (PCA on sentence trials, decoding image trials) was then performed. Tables 5 and 6 display the Wilcoxon statistics for the sequentially incoherent/coherent distinction in this cross-modal validation, for the first, and second stimuli, respectively. Recall that when the first image or sentence is presented, the subject cannot yet determine if it is coherent or incoherent, and so we predict no decoding can be made on the data from stimulus 1. Indeed, this is confirmed in Table 5. No coherent vs incoherent differences are significant for stimulus 1. We see that in Table 6, the data for stimulus 2 reveals a different story. When the PCA was performed on the image data, the first component reliably distinguished between sequential coherence for images, in the two late periods. More importantly, this component also reliably made the coherence distinction for the sentence trials, in both late periods. In the complementary sense, PCA performed on sentence data could then reliably make the coherence distinction both for sentences and images. This argues that the processing of sequential coherence has common neural substrates for images and sentences.

**Table 5.** Sequential coherence cannot be discriminated for Stimulus 1. Control test of PCA decoding of coherence of the first stimulus, depending on training and testing conditions. Wilcoxon p values for Sequentially Coherent - Incoherent differences for stimulus 1. Significant values (p < 0.05) indicate that the coherence difference is reliable. Upper row provides results from training on Image data, lower row from training on Sentence data. Parentheses () indicate the crucial transfer conditions where training is in one modality, and testing in the other. Here we expect no differences to be significant, since the coherence distinction cannot be made at the first stimulus.

|  | **Late 1 (750-1000)** |  | **Late 2 (1600 – 1800)** |  |
| --- | --- | --- | --- | --- |
|  | **Test on Image1** | **Test on Sentence1** | **Test on Image1** | **Test on Sentence1** |
| **Train on Images** | 0.616 | (0.845) | 0.647 | (0.145) |
| **Train on Sentences** | (0.586) | 0.777 | (0.616) | 0.144 |

**Table 6.** Sequential coherence is cross-modally decoded for Stimulus 2. Transfer test of PCA decoding of sequential coherence of the second stimulus, depending on training and testing conditions. Wilcoxon p values for NonSEQ – SEQ differences for stimulus 2. Significant values (p < 0.05) indicate that the Sequential Coherence difference is reliable. Upper row provides results from training on Image data, lower row from training on Sentence data. Parentheses () indicate the crucial transfer conditions where training is in one modality, and testing in the other. Here we expect all differences to be significant.

|  | **Early (750-1000)** |  | **Late (1600-1800)** |  |
| --- | --- | --- | --- | --- |
|  | **Test on Image2** | **Test on Sentence2** | **Test on Image2** | **Test on Sentence2** |
| **Train on Images** | 0.016 | (0.048) | 0.031 | (0.009) |
| **Train on Sentences** | (0.020) | 0.043 | (0.039) | 0.009 |

## Discussion

In the current research we analyze neural activity generated while subjects are exposed to complex human-activity-related stimuli presented in two-event narrative sequences. The sequences are presented in two distinct perceptual formats or dimensions: sentences and images. Our objective is to test the hypothesis that common neural processes are involved in making sense of these narrative sequences in the sentence and image conditions. We previously performed a related experiment where we recorded brain activation using fMRI while subjects were involved in understanding single visual images and written sentences. This revealed a network of brain regions common to processing sentences and images suggesting that the understanding process involves mapping the understood situation into one’s own experiential representation of meaning (Jouen et al 2015). This is coherent with the semantic network identified by Binder et al (2009), and extends this into the visual image domain. Further evidence for a common semantic representation for words and pictures has been provided by cross-modal decoding of object categories from the fMRI signal (Shinkareva et al 2011). Here we examine the neural dynamics associated with engaging the semantic system as required for understanding two-element narratives made up of images or sentences. We orient the discussion around the principal results, particularly the semantic processing associated with the late EEG responses, the corresponding behavior, and steps towards conceptual and neural implementations of explanatory models.

### Early Specific & Common Semantic Effects

Already by 300 ms. we observe two effects in the EEG signal – a modality effect with distinct processing for sentences and images, and more interestingly, for both sentence and image modalities we also observe common processing revealed as the sequential coherence effect. Such early responses have been observed in language processing for syntactic class expectation violations (Neville et al 1991), and in image processing in response to comic strip panels in which the motion lines were in conflict with the depicted motion, or when they were absent (Cohn & Maher 2015). This suggests that this early positivity is a response to a violation of an expectation with respect to the narrative structure linking the two successive stimuli.

Interestingly we observed a negativity that can be compared to the N400 in response to the incoherent images, but not sentences. Cohn et al (2012) similarly observed N400 effects in response to image sequences that violated narrative structure and semantic structure, and Paczynski and Kuperberg (2012) note different combinations of stimuli can produce late positive responses with and without the N400. This is similar to the discourse effects observed by van Berkum and Hagoort (Hagoort & van Berkum 2007, van Berkum et al 2003, Van Berkum et al 2005, Van Berkum et al 1999), where an N400 can be evoked by an otherwise neutral sentence that has been rendered anomalous by preceding context.

### Late Common Semantic Processing

The main result is the reliable observation of late centrally distributed positive ERP responses during the processing of sentences and images in the sequentially incoherent condition. The late positivity that we observe in the 750-1000 ms timeframe is similar to those that have been observed when sentences and images do not fit with the previously established context (Bornkessel-Schlesewsky & Schlesewsky 2008, Brouwer et al 2012, Kuperberg 2007, Xiang & Kuperberg 2015). While it is beyond the scope of the current analysis to retrace the details of the evolution of the P600 from its syntactic processing origins (Osterhout & Holcomb 1992) to its more recent manifestation in the complex world at the interface of syntax and semantics (see below), we can attempt to situate our finding of a late frontal positivity that is associated with a form of making meaning both for text and visual narratives.

Thus, while frontal positivities were initially produced in response to syntactic anomalies, they have since been identified in a number of situations that do not involve syntactic recovery. For example Kaan and Swaab (2003) observe a late (500-900 ms) frontal positivity associated with ambiguity resolution and/or increases in discourse complexity. Kuperberg (2007) reviews studies where late positivities are evoked, with and without syntactical anomalous sentences. Of particular interest, in the sentence “Every morning at breakfast the <u>eggs</u> would eat …”, the response to eggs (which is semantically anomalous as inanimate eggs cannot eat) was a robust P600, in the absence of an N400 (Kuperberg et al 2003). Kuperberg suggests that some degree of semantic association between a verb and its arguments may trigger a P600. Similarly, Xiang and Kuperberg (2015) have argued that a late posterior positivity component is triggered when a near certain prediction is followed by an input that requires a switch to a new generative model representing relationships between events. Paczynski and Kuperberg (2012) advocate the P600 as reflecting a conflict between semantic memory-based predictions, and the detection of propositional incoherence. Brouwer et al (2012) consider that late positivities are invoked by semantic integration processes. Our sequentially incoherent condition would thus invoke such integration processes producing a prediction error.

Accordingly, in the same time frame, we observed increases of near-gamma band activity (35-45 Hz) for sequentially coherent stimuli, both for sentence and image processing. Such increases in gamma band activity have been proposed to reflect integrative processing through enhanced coherence of neural activity in multiple interacting regions (Schneider et al 2008, Senkowski et al 2008). Thus, in the sequentially coherent conditions, participants have established a context with the first stimulus and can then proceed with further integration and processing with the second stimulus when it is sequentially coherent. This is associated with increased gamma activity. Gamma activity is more generally associated with a number of higher cognitive functions including controlling and organizing memory traces (Keizer et al 2010), and semantic processing in sentence comprehension (Hagoort et al 2004, Hald et al 2006).

### Cross-modality decoding

The ERP and time frequency analyses revealed late processes that were common to sentence and image processing, suggesting a common underlying neural mechanism. These findings based on ERP and time-frequency analyses are reinforced by a novel approach using PCA and cross validation over sentences and image. If the neural dynamics are related for processing the sequentially incoherent stimuli for sentences and images, then a pattern decoder trained on sentences should be able to decode sequential coherence in images, and vice versa. To investigate this we employed multivariate analysis (principle component analysis – PCA) for decoding semantic coherence across modalities. The principal component analysis revealed that the EEG signal reliably encodes information about the modality of the stimuli (sentence vs image), and the status of the second stimulus as being sequentially coherent or incoherent with respect to the first. As a multivariate analysis tool, PCA allows the opportunity of generating principal components from one set of data, and then testing another set of data using those components in order to determine whether the second data set is related to the first (Brouwer & Heeger 2009). Analyses were thus performed to determine whether the principal components that characterize narrative coherence in sentence processing would allow decoding for image processing, and vice versa. The cross-validation results revealed that indeed this was the case. When PCA was applied to sentences, the first component reliably distinguished coherent vs. incoherent sentences and images, and only for the crucial second stimulus of the pair. The same positive outcome was observed when applying the complementary analysis, i.e. running PCA on the image data and testing on the sentence data. Thus, this cross-modal generalization argues that the same processes are at work in narrative integration of the successive stimuli for sentences and images. We can suggest that this type of situation can be established in a narrative context, so that while the second sentence contains no inherent violation, it is anomalous with respect to the narrative context established by the first sentence.

While the actual cognitive processing that is associated with these late EEG patterns revealed by ERP, time-frequency and PCA analyses remains to be clarified, we can conclude that sequential incoherence in short narratives reliably evokes late positivities with similar timing and topography, as well as similar TF patterns and PCA signatures for sentences and images. This common electrophysiological profile is evidence for common underlying neurophysiological processing which is further supported by behavioral observations. One might consider that the responses observed for incoherent stimuli are simply related to detection of an unpredicted stimulus. The extended processing differences during the later periods are not consistent with a simple surprise response (Brouwer et al 2012, Sitnikova et al 2008), and suggest additional processing. The cross-modal decoding argues further in this sense. The underlying neural processing of surprise must be at a narrative semantic level, which implies that to be surprised requires a certain level of understanding that is independent of the stimulus modality, which is consistent with our hypothesis.

### Behavioral Correlates of Late Semantic Processing

This leads to the question of what it is that subjects are actually doing in our task. One suggestion is that without any intention or will to do so, subjects naturally engage with the stimuli and attempt to make sense of them, to connect them. Data from studies of embodied language indicate that this is the case for language processing (Aziz-Zadeh et al 2006), and we can consider that it is similar for image processing. In separate studies we have begun to behaviorally investigate this integrative process in the domain of sentence processing (Madden-Lombardi et al 2015). There, we exposed subjects to successive sentences in pairs that were either sequentially coherent or not. Subjects found the second sentence in the sequentially incoherent pairs less easy to mentally imagine than those in the sequentially coherent condition, and they also required additional time to make the judgement (Madden-Lombardi et al 2015). This suggests that in the sequentially coherent pairs, the representation of the second sentence is already (partially) included in the representation evoked by the first sentences, whereas in the sequentially incoherent pairs, additional processing is required. This additional processing may be reflected in the late positivities we observe in the present study, while the integration of the coherent image may be reflected by the increase near gamma response. This processing may be part of attempting to make sense in the context of narrative integration.

According to the event indexing model of Zwaan and colleagues (Zwaan 1999, Zwaan et al 1995) coherence can be measured along five dimensions: time, space, causation, motivation and protagonist. In the sequentially incoherent conditions, while relations in the dimensions of time and space may be broken, the protagonist relation remains, so that the sequentially incoherent conditions can be considered as a shift in time or storyline rather than a completely incoherent or unrelated event (Madden-Lombardi et al 2015).

### Towards Models of Neural Dynamics of Semantic Structure Processing

How can such semantic structure processing be mapped onto neural dynamics? In a long-term effort to respond to this question, and based on the property of massive local recurrent connections in primate cortex (Goldman-Rakic 1987) we have developed a class of recurrent network models that are sensitive to sequential structure: from the serial and temporal structure of sensorimotor sequences (Dominey 1998a, Dominey 1998b), to abstract rule structure in language and artificial grammars (Dominey 2005, Dominey et al 2009, Hinaut & Dominey 2011, Hinaut & Dominey 2013). These models can learn and generalize the mapping from grammatical structure to meaning, and provide explanations for certain human language-related neurophysiology (Hinaut & Dominey 2013). Based on structural regularities in the sentence-meaning data used during training, the recurrent network model can anticipate the next words in an incoming sentence, and when low probability words arrive, the model displays a perturbation of its activity due to adjustment between the anticipatory activity that is inappropriate to what is actually perceived. These perturbations are related to the P600 that is observed in response to sentence structure violations (Hinaut & Dominey 2013).

We have recently extended the model of grammatical construction to that of narrative construction. By analogy, where a grammatical construction defines the relation between words in a sentence and their relation to arguments in a predicate-argument event representation, a narrative construction defines the structural relations between sentences in a narrative and multiple interrelated events and relations between them in a situation model (Mealier et al 2017). Exploiting this analogy, at the level of the narrative construction, we would predict that when incoming sentences that have low probability (with respect to narrative structures learned in the training corpus) arrive, they will likewise result in a perturbation of the model’s prediction corresponding to a late positivity.

This suggests the existence of generalized structure processing mechanisms whose response to unpredicted structure – including semantic structure - will be a late positivity (Brouwer et al 2012, Xiang & Kuperberg 2015) Here we begin to accumulate evidence that such structural processing may apply as well at the level of narrative structure that can be evoked by processing sentences or images that depict unfolding human events.

## Conclusion

The groundbreaking work of Vandenberghe et al (1996), provided evidence for a common semantic system for words and pictures. This was further confirmed at the semantic category level by cross-modality decoding of semantic categories (Shinkareva et al 2011). Jouen et al (2015) extended this line of research, revealing a broadly extended semantic system common to the representation of the semantics of human event evoked by sentences and complex images, with extended anatomical and functional connectivity (Jouen et al 2018). The current research reveals evidence for common neural dynamics for semantic processing of short narrative sequences made up of sentences or images. Using EEG we observe common spatiotemporal patterns of brain activity when subjects encounter sequentially incoherent stimuli in the image or sentence modalities. This is revealed in the ERP domain as a set of late central positivities, and in the time-frequency domain as an increase in near-gamma power. This is coherent with related research examining sentence and image processing separately. Importantly, in the stimuli that we use to asses this sequential narrative processing, there is nothing inherently wrong with the second stimulus in these sequentially incoherent pairs: it is only their relation with the first stimulus that is manipulated in order to produce a sequential incoherence.

Further supporting the hypothesis of common modality-independent processing, multivariate analysis discovers this sequential coherence effect in one modality, and can then be used to decode sequential coherence in the other modality, arguing for a common neurodynamics of meaning integration, independent of the input modality. This demonstrates that the brain tracks coherence not only within individual sentences and images, but across multiple stimuli in a narrative context, and that this mechanism is at least partially independent of the stimulus modality. The question of the nature of the contents of these representations remains a topic of future research. A growing body of research now addresses the representation and decoding of the contents of meaning, e.g. (Shinkareva et al 2011, Wehbe et al 2014). Future research should attempt to apply these techniques across extended narrative in sentence and visual modalities in the further characterization of the common semantic system.

## Acknowledgements

This research was supported by funding from the French ANR Comprendre, and from the European Community through grants FP7-ICT 231267 Project Organic, FP7-ICT-270490 Project EFAA, and FP7-ICT-612139 Project WYSIWYD.

